# Q^INT^, a novel simplified QF/QUAS expression system with integrated temporal expression control

**DOI:** 10.64898/2026.04.16.719020

**Authors:** Ji-Eun Ahn, Hubert Amrein

## Abstract

Binary gene expression systems have played a major role in uncovering the function of genes, cells and organ systems during and after development. Employed initially in model systems such as flies and mice, advances in gene technology have vastly expanded the number of species in which these systems can be deployed. One of their limitations is the challenge of imposing temporal expression control. Here, we report the incorporation of temperature-sensitive intein modules with different temperature profiles into QF2, the transcription factor anchoring the Q-system widely used in *Drosophila* and introduced recently into several insect species and the zebrafish *D. rerio*. Intein removal from temperature-sensitive QF2_INT^ts^ activators in *Drosophila* larvae and adult flies raised under permissive conditions (18°C to 25°C) renders QF2 active and drives strong expression of *QUAS* reporters. In contrast, raising *Drosophila* at restrictive temperatures (25°C to 30°C) keeps QF2_INT^ts^ in a non-functional state unable to bind DNA and therefore keeping QUAS reporters inactive. We further show that reporter expression can be turned on and off both in larvae and flies by changing temperature conditions between restrictive to permissive. Finally, expression of the inward rectifying potassium channel (*QUAS-Kir2.1*) under the control of QF2_INT^ts^ drivers in olfactory neurons and pan-neuronally was effectively suppressed in flies raised under restrictive temperature conditions with no impact on olfactory acuity or survival, respectively. In contrast, when respective flies were shifted to permissive temperature-condition, flies lost olfactory acuity and became paralyzed within one to four days, respectively. Thus, the intein-based Q-system, henceforth referred to as Q^INT^ system, renders the difficult to employ drug-dependent suppressor QS unnecessary, greatly facilitating temporal regulation of gene expression. Having several advantages over the GAL4 system, the Qsystem has gained broad acceptance in other insect species and the zebrafish *D. rerio*, and thus the temperature-regulatable Q^INT^ system can be implemented in many other organisms, including disease-transmitting mosquitoes and poikilotherm vertebrate animals to study gene and cell function at any time during or after development.

## Introduction

Binary expression systems are powerful genetic tools for manipulating gene function with cell and tissue specificity. Three of these systems – the GAL4 (Brand and Perrimon 1993), LexA (Lai and Lee 2006) and Q-systems (Potter et al. 2010; Potter and Luo 2011) - have been developed and refined to different extents in the versatile *Drosophila* model system, providing fundamental insights into gene function across a wide array of different cell types during development and in adult flies. All these systems are based on the same basic principle: a driver composed of a promoter imposing gene-specific expression on a transcriptional activator (GAL4, LexA or QF) and a reporter gene of interest cloned downstream of the respective activator- responsive regulatory sequence (*UAS*, *lexA_op_*, or *QUAS*). Gene expression can be further refined using repressors such as GAL80 (for GAL4, Ma and Ptashne 1987) or QS (for QF, Huiet and Giles 1986; Giles et al. 1991). These suppressors bind to their cognate activators, thereby blocking transcriptional activation by preventing engagement with the basic transcription machinery. These suppressors are regulatable temporally, either by ambient temperature conditions in the case of GAL80^ts^, a temperature-sensitive GAL80 variant that is inactive above 30 °C (McGuire et al. 2003), or the presence of dietarily provided quinic acid that binds QS and prevents its interaction with QF (Potter et al. 2010).

With the development of genome editing tools, such as CRISPR/Cas9 and TALENs (Liu et al. 2012; Zirin et al. 2022; Kanca et al. 2022; Zirin et al. 2024), binary expression systems can now be deployed to virtually any organism. Early studies introduced the GAL4 system into mosquito species, specifically the yellow fever mosquito *Aedes aegypti* (Kokoza and Raikhel 2011; B. Zhao et al. 2014) the malaria mosquitoes *Anopheles stephensi* (O’Brochta et al. 2012) and *Anopheles gambiae* (Lynd and Lycett 2012). However, effective and reliable implementation of GAL4, especially when expressed under the control of strong ubiquitously active or pan-neural promoters remained elusive for many genes, due to cell toxicity (Coutinho-Abreu and Akbari 2023; Z. Zhao et al. 2021) In zebrafish, the other organism into which the GAL4 system was introduced, its effective use is challenging because of CpG sequences present in the UAS binding site (CGG– N_11_–CCG), which are subject to DNA methylation and can lead to silencing of reporter expression (Goll et al. 2009; Akitake et al. 2011). Because of these limitations, investigators have introduced the Q-system into both mosquitoes and zebrafish. Although QF can exhibit cellular toxicity similar to GAL4, this issue has been substantially reduced through the development of second and third generation activators, such as QF2 and QF2^w^. Furthermore, transgene silencing is a not a concern in the Q system (Subedi et al. 2014; W. Huang et al. 2014) because the QUAS regulatory sequence is devoid of CpG sites.

The main drawback of the Q-system is the deployment of temporal control via its repressor QS. In contrast to GAL80^ts^, whose function for temporal suppression of GAL4 is reversibly controlled by shifting animals from permissive (19°C) to restrictive (30°C) temperature and vice versa, QS activity is dependent on the small polyhydroxy cyclic alcohol quinic acid. Specifically, QS binding to and suppression of QF/QF2/QF2^w^ can only be relieved in the presence of quinic acid, which must be provided through the diet. However, application of quinic acid presents several challenges, including its sour and slightly bitter taste, low efficacy in crossing the blood brain barrier, overall toxicity in some animals leading to tissue and organ defects when applied during development (especially in fish), and slow uptake (due to poor feeding) and clearance (Potter et al. 2010; Wei et al. 2012; Ghosh and Halpern 2016; Riabinina et al. 2015; Riabinina and Potter 2016).

Here, we present a strategy for achieving temporal control of the Q-system without the need of an additional suppressor component by incorporating temperature-dependent intein modules into QF2. Inteins are self-splicing protein elements encoded within a small number of genes of many prokaryotes and some unicellular eukaryotes, where their excision restores the function of the host proteins.

We re-generated previously characterized, temperature-sensitive inteins (INT^ts^) derived from the *Saccharomyces cerevisiae VMA1* gene and inserted them into QF2 to create QF2_INT^ts^ drivers that can be activated and silenced by shifting *Drosophila* larvae or flies from restrictive to permissive temperature and vice versa. These constructs were placed under the control of the *Drosophila olfactory receptor co-receptor* (*Orco*) and the *neuronal Synaptobrevin* (*nSyb*) promoters to enable cell-type-specific, temperature-gated activation of QF2. We show that functional expression of *Orco-QF2_INT^ts^* and *nSyb-QF2_INT^ts^* in *Drosophila* is temperature dependent. Moreover, we find that reporter expression in flies and larvae raised at permissive temperature is comparable to that when driven by standard (intein-less) *Orco-QF2* and *nSyb*-*QF2* drivers. Furthermore, down-shift and up-shift experiments show that reporter expression can be turned on and off, respectively, indicating that intein-splicing too can be switched on and off by simply lowering or increasing the ambient temperature. Finally, we show that *nSyb/Orco*- *QF2_INT^ts^; QUAS-Kir2.1* animals raised at restrictive temperature are rescued from embryonic lethality or loss or olfactory acuity, phenotypes observed at permissive temperature or when *QUAS-Kir2.1* is combined with a standard *nSyb/Orco*-*QF2* driver. These observations eastablish that QF2_INT^ts^ activators can effectively be used to avoid severe phenotypic effects of broadly expressed reporters that affect essential cellular functions. Since the Q-system has been widely adapted for other invertebrate systems, as well as zebrafish *D. rerio*, temporal gene expression control should now also become feasible for many other poikilotherm animals.

## Results

### Engineering QF2_INT activators

Temperature-dependent splicing of VMA1 inteins from foreign host proteins, specifically GAL4 and GAL80, was previously demonstrated in *Drosophila* (Zeidler et al. 2004). Given the broader species pool for the Q system when compared to the GAL4 system, we integrated temperature-sensitive VMA1 intein modules tested in yeast (Tan et al. 2009) into QF2, thereby bypassing the need of the drug-dependent QS. We first tested the efficacy of intein splicing from QF2 protein using the wild-type (temperature-insensitive) intein inserted upstream of four different cysteine residues, residing in the zinc finger DNA binding motif (C79, C86 and C103) or the C-terminal activation domain (C340; Figure 1A). While a cysteine residue downstream of the intein is the only strict requirement for VMA1 intein self-splicing, both amino acid context near the insertion site, as well as secondary structure of the host protein can contribute to splicing efficacy (Apgar et al. 2012). Thus, insertion into the zinc finger motif, which is expected to disrupt QF2 binding to DNA, will allow conclusive assessment for VMA1 intein self-splicing. Western blot analysis of GAL4 containing VMA1 inteins results in expected size reduction of the host protein to its original size, suggesting precise intein removal via self-splicing (Zeidler et al. 2004; Tan et al. 2009). However, lack of a QF2 antibody does not offer any means to check for precise removal of the intein in this study.

**Figure 1.**
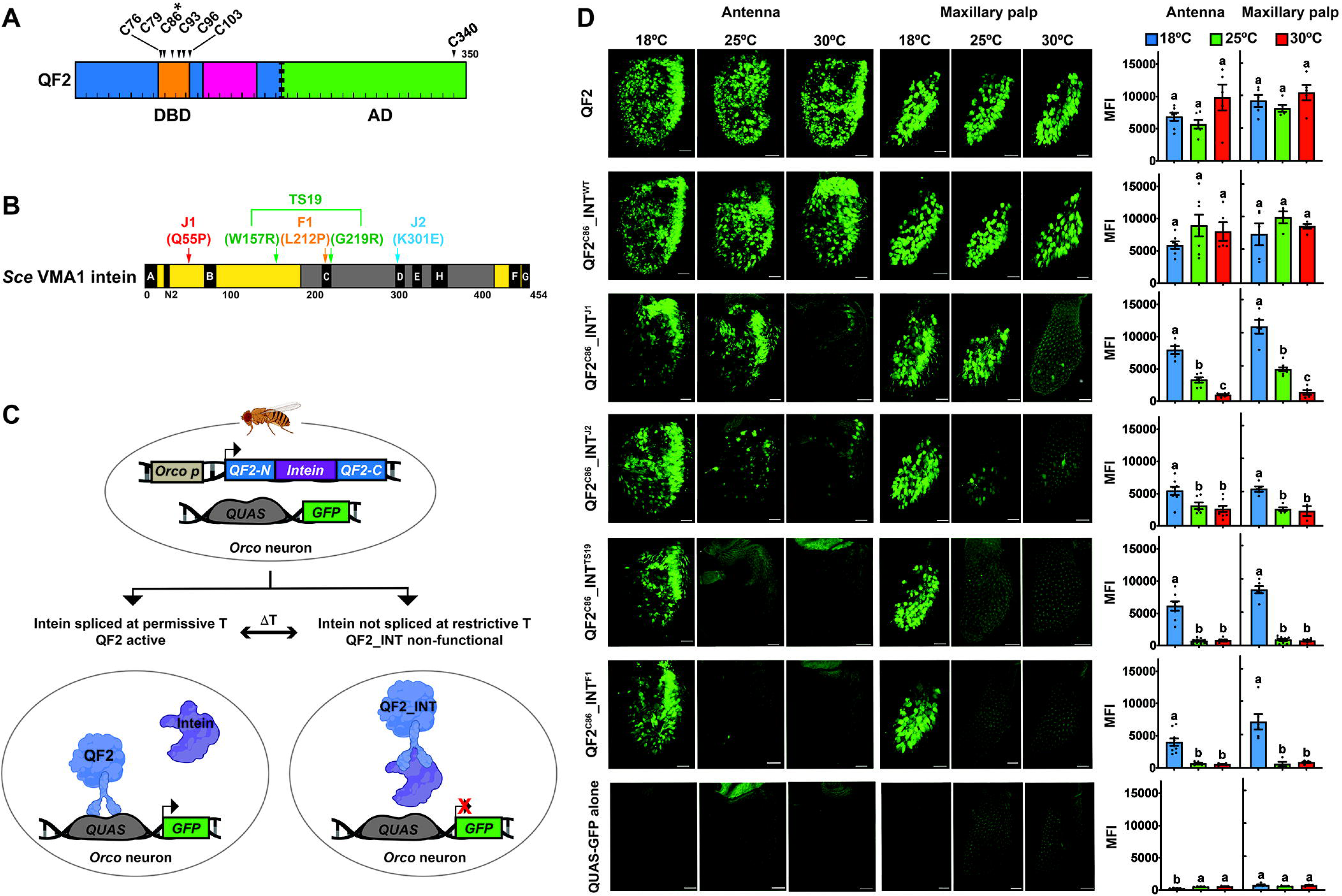
Intein-containing, temperature-sensitive QF2 activator. **A)** Structure of QF2 activator. DBD: DNA-binding domain (blue) and AD: activation domain (green). Zn_2_/Cys_6_ zinc finger motif and dimerization domain are shown in orange and red, respectively. QF2 with intein inserted at C86 (QF2^C86^_INT) were used throughout this study. The dashed line indicates the location of the middle domain removed in the original QF to avoid toxicity caused by broad expression of QF. **B)** Structure of the *S. cerevisiae* (*Sce*) VMA1 intein. Splicing domains and homing endonuclease (HEN) domain in yellow and grey, respectively. Conserved motifs essential for protein splicing (A, N2, B, F and G), and endonuclease activity (C, D, E and H) are indicated as black boxes. Positions of temperature-sensitive mutations used in this study are indicated. **C)** Intein-splicing dependent Q^INT^-system: Temperature-sensitive inteins were inserted at C86 of the QF2 coding sequence. Self-splicing of the intein at permissive temperature connects the two QF2 fragments and reconstitute a functional QF2 protein, while at restrictive temperature, the intein is retained within the DNA binding domain, abolishing QF2 activity. *Orco* or *nSyb* promotors are used to express QF2_INT^ts^ drivers. At permissive temperatures, reconstituted functional QF2 enables QUAS-dependent GFP expression, while at restrictive temperatures, splicing is impaired leaving intein module within QF2, and disabled QF2 fails to activate QUAS-GFP reporters. **(D)** QF2^C86^_INT^ts^ enables temperature-dependent GFP expression in *Orco* neurons. Confocal images of flies containing indicated *QF2* drivers and *10xQUAS-6xGFP* reporter show live GFP expression at indicated temperatures (18°C, 25°C or 30°C). QUAS-GFP alone serves as a control for leaky expression. Fluorescence imaging of GFP was performed on whole-mount preparations of antennae and maxillary palps from flies of the following genotypes: *w^1118^;10xQUAS- 6xGFP/+;Orco-QF2/+*, *w^1118^;10xQUAS-6xGFP/+;Orco-QF2^C86^_INT^WT^/+*, *w^1118^;10xQUAS- 6xGFP/+;Orco-QF2^C86^_INT^J1^/+*, *w^1118^;10xQUAS-6xGFP/Orco-QF2^C86^_INT^J2^*, *w^1118^;10xQUAS- 6xGFP/+;Orco-QF2^C86^_INT^TS19^/+*, *w^1118^;10xQUAS-6xGFP/+;Orco-QF2^C86^_INT^F1^/+* and *w^1118^;10xQUAS-6xGFP/+*. GFP expression was quantified and displayed as MFI (mean fluorescence intensity) in the histogram. Each bar represents the mean ± SEM of MFI (n = 3-11 MFI measurements in the antenna and maxillary palp). Dots indicate MFI measured in individual antenna and maxillary palp. Ordinary one-way ANOVA with Tukey’s multiple comparison tests (p < 0.05). Different letters indicate statistically significant differences. Note that enhanced appearance of hairs, especially in images of maxillary palps is due to autofluorescence, and not GFP expression. Scale bars are 20 µm.

We generated several independent driver lines with each of *QF2_INT^WT^* modules under the control of *Orco* promotor (Figure 1B and 1C). *Orco* which encodes the core subunit of OR based tetrameric olfactory receptor complexes, is broadly expressed in all olfactory sensory neurons (OSNs) of basiconic sensilla, which comprise about 75% of all antennal OSNs and all OSN of the maxillary palps (Larsson et al. 2004). We crossed these drivers to a 10x*QUAS-6xGFP* reporter strains and assessed GFP expression in the antenna and maxillary palp (Supplementary Figure 1). None of the QF2 driver lines with intein insertions at C79 or C103 led to GFP expression, suggesting that self-splicing does not occur or is inefficient from these locations, presumably due to structural constraints (Supplementary Figure 1). In contrast, all intein containing QF2 drivers with intein insertions at C86 or C340 showed robust GFP expression in OSNs of both the antenna and the maxillary palp, an expression indistinguishable from that observed with the standard *Orco-QF2* driver (Supplementary Figure 1). We next generated four *Orco-QF2_INT^ts^* drivers containing temperature-sensitive inteins with single or double amino acid substitutions previously associated with temperature-dependent splicing profiles (Tan et al. 2009) and inserted them into the splicing competent C86 location of QF2, resulting in *Orco- QF2^C86^_INT^J1^*, *Orco*-*QF2^C86^_INT^J2^*, *Orco-QF2^C86^_INT^TS19^* and *Orco-QF2^C86^_INT^F1^* (Figure 1B-1C and Supplementary Table 1). We then analyzed expression in OSNs of such flies also harboring the *10xQUAS-6xGFP* reporter at 18^°^C, 25^°^C and 30^°^C, along with two positive controls (standard *Orco-QF2*, *Orco-QF2_INT^WT^*) and a negative (reporter-only) control (Figure 1D). *Orco-QF2* and *Orco-QF2^C86^_INT^WT^* showed robust *GFP* expression in OSNs of both the antenna and the maxillary palps under all three temperature conditions, while the reporter line alone lacked any expression. Three drivers showed no (*Orco-QF2^C86^_INT^TS19^* and *Orco-QF2^C86^_INT^F1^*) or rare, sporadic expression at 30°C (*Orco-QF2^C86^_INT^J1^*), while *Orco-QF2^C86^_INT^J2^* exhibited weak, but consistent expression in about 15 to 20 antennal and 3 to 5 maxillary palp neurons (corresponding to about 1.5 to 2% of OSNs). Importantly, all four lines showed robust expression at 18°C (Figure 1D) mimicking that of *Orco-QF2* and *Orco-QF2^C86^_INT^WT^*. At 25^°^C, *Orco-QF2^C86^_INT^J1^* provided robust expression and *Orco-QF2^C86^_INT^J2^* showed expression in a fraction of OSNs of olfactory organs, while in *Orco-QF2^C86^_INT^TS19^* and *Orco-QF2^C86^_INT^F1^* expression was entirely absent at this temperature. Lastly, we conducted quantitative fluorescent imaging experiments of antenna and maxillary palps dissected from flies of these genotypes but subjected to immunohistochemistry and anti-GFP antibody staining (Supplementary Figure 2), which aligned completely with the observation made using quantitative live fluorescent imaging (Figure 1D).

These findings confirm that intein splicing of all four temperature-sensitive modules is temperature-dependent. However, while the temperature profiles of *QF2^C86^_INT^F1^* in flies was similar to that reported previously in the context of GAL4 when expressed in yeast (Tan et al. 2009), *QF2^C86^_INT^TS19^* was only expressed at the lowest temperature, while in yeast, this module was robustly spliced out from GAL4 up to 25 °C (Supplementary Table 1). Surprisingly, the two single mutations in *Orco-QF2^C86^_INT^J1^* (Q55P) and *Orco-QF2^C86^_INT^J2^* (K301E) exhibited a similar temperature profile as the same mutation (S40 and S7) in the INT^TS19^ background (W157R/ G219R) when spliced from GAL4 (Tan et al. 2009; Supplementary Table 1). In summary, the four different temperature-sensitive inteins exhibit distinct, reproducible splicing from QF2 across a range of temperature when expressed in *Drosophila* OSNs.

### Reversible, temperature-dependent control of *QF2_INT^ts^*

To determine whether QF2_INT^ts^ drivers can be switched on and off, we subjected *Orco- QF2^C86^_INT^J1^; 10xQUAS-6xGFP* flies to temperature-shift paradigms (Figure 2A). One day old *Drosophila* flies raised at 25°C were shifted to the restrictive temperatures (30°C) for 6 days, followed by a down shift to either 25°C or 18°C for a total of 4 days. GFP expression in most if not all OSNs was strong following development at 25 °C but mostly lost after 3 days and completely absent after 6 days at 30°C (Figure 2B). When these flies were downshifted to 25°C, GFP expression was recovered in a few OSNs within a day and slightly increased in maxillary palp OSNs after an additional 3 days. When flies were downshifted from 30°C to 18°C, GFP positive neurons were more abundant in both organs after 1 day compared to downshift to 25°C, while after four days at 18°C, expression was restored in most if not all OSNs of both olfactory organs. Quantification of GFP expression confirmed that expression was fully recovered when flies were shifted within 1 to 4 days from the restrictive temperature (30°C) to 18°C (Figure 2C). Together, these observations show that QF2^C86^_INT^J1^ function can be switched on and off by lowering and raising temperature from restrictive to permissive conditions, and vice versa.

**Figure 2.**
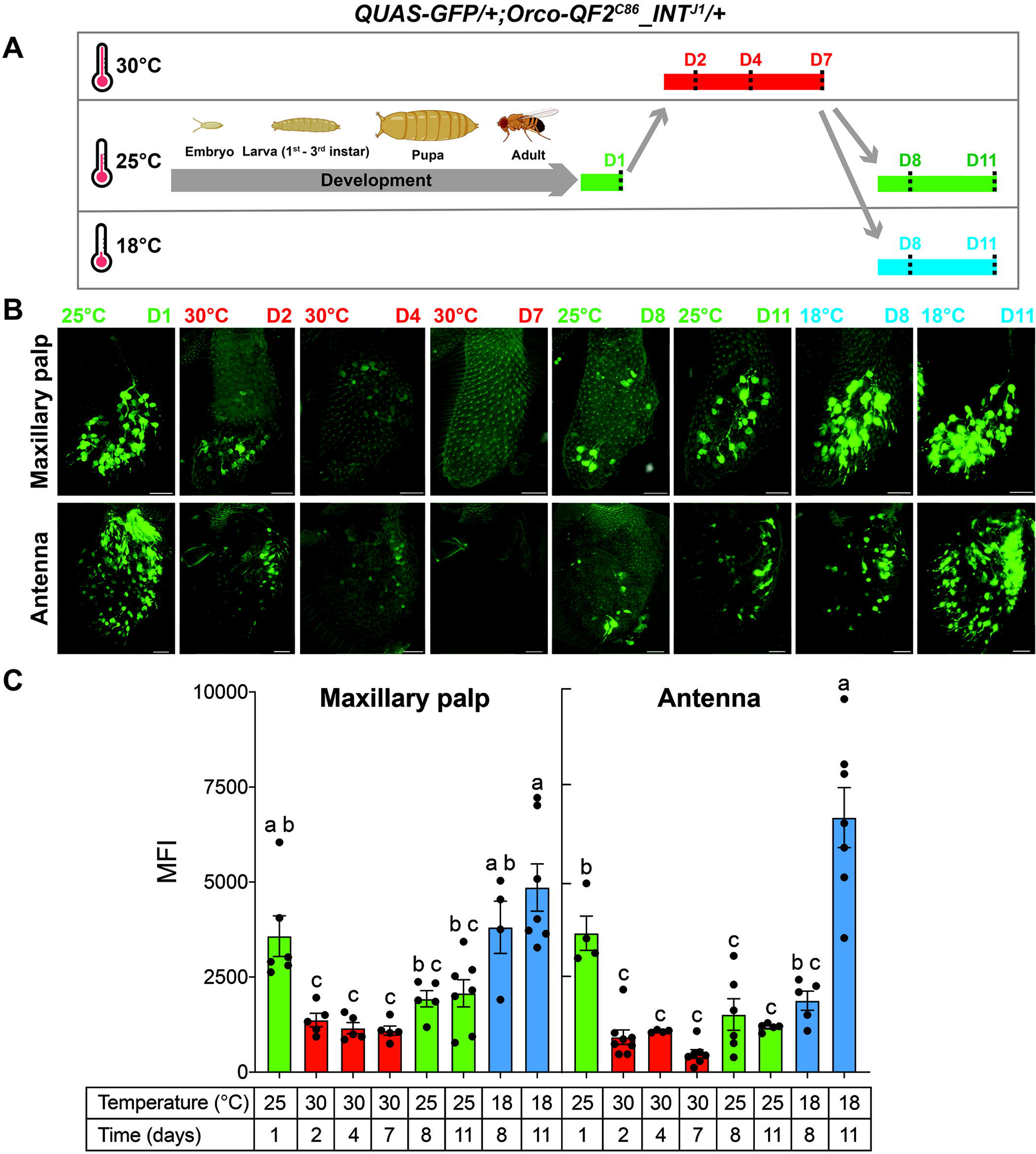
Reversibility of QF2_INT^J1^ activity in *Orco* neurons using temperature-shifts. **(A)** Experimental temperature-shift paradigm. *w^1118^;10xQUAS-6xGFP/+;Orco-QF2^C86^_INT^J1^/+* flies were raised at 25 °C throughout development and subjected to temperature shifts between permissive (18°C/25°C), and restrictive (30°C) conditions for the indicated durations. **(B)** Temperature-dependent control of QF2-intein is reversible and tunable in *Orco* neurons. Confocal images of live GFP expression in maxillary palps (top row) and antennae (bottom row) at each time point. GFP is efficiently suppressed after prolonged exposure to restrictive temperature (30°C, D2-D7) and progressively restored upon shifting back to permissive temperatures (25°C and 18°C). Note that recovery of expression resumes faster at lower permissive temperature. Scale bars are 20 µm. **(C)** Quantification of GFP expression driven by QF2^C86^_INT^J1^ in *Orco* neurons. Each bar represents the mean ± SEM of MFI (mean fluorescence intensity) (n = 4-8 MFI measurements in the antenna and maxillary palp). Dots indicate MFI measured in individual antenna and maxillary palp. Ordinary one-way ANOVA with Tukey’s multiple comparison tests (p < 0.05). Different letters indicate statistically significant differences.

### *nSyb-QF2_INT^TS19^* provides temporal pan-neural control in larvae

To generate a regulatable activator that is more broadly expressed in the nervous system, we cloned QF2^C86^_INT^TS19^ and QF2^C86^_INT^WT^ downstream of the *neuronal Synaptobrevin* (*nSyb*) promoter and generated several new *nSyb-QF2^C86^_INT^TS19^* driver lines. We selected the *nSyb* promoter, because it is expressed in all neurons (Riabinina et al. 2015), and we chose INT^TS19^ for these experiments, because this module has a lower temperature on-off range than INT^J1^ and INT^J2^, providing a larger time window for temperature shift experiments (larvae growth and development is half as fast at 18°C compared to 25°C). Indeed, *nSyb-QF2^C86^_INT^WT^*;*10xQUAS- 6xGFP* larvae showed robust GFP expression in the brain and ventral nerve cord (VNC), as well as dispersed peripheral sensory neurons, at all temperatures (18°C, 25°C, and 30°C) (Figure 3A, middle panels). By contrast, larvae expressing *nSyb-QF2^C86^_INT^TS19^* exhibited the expected temperature dependence observed in OSNs under the control of the *Orco* promoter (Figure 1D). Specifically, at 18°C, GFP expression replicates that of *nSyb-QF2^C86^_INT^WT^*, with robust labeling in the brain, VNC and sensory neurons (Figure 3A, right panels). In contrast, at 25°C and 30°C, no GFP expression is visible, indicating that QF2 remains in its non-spliced intein-containing form that is unable to bind QUAS and activate reporter gene expression. To test how quickly intein splicing can be induced, we downshifted first instar larvae from the restrictive (25 °C) to the permissive (18°C) temperature and examined and quantified GFP protein levels at 6-hour intervals (Figure 3B). These experiments demonstrate that rapidly induced intein splicing results in reporter expression within 6 hours, reaching steady state level within 24 hours.

**Figure 3.**
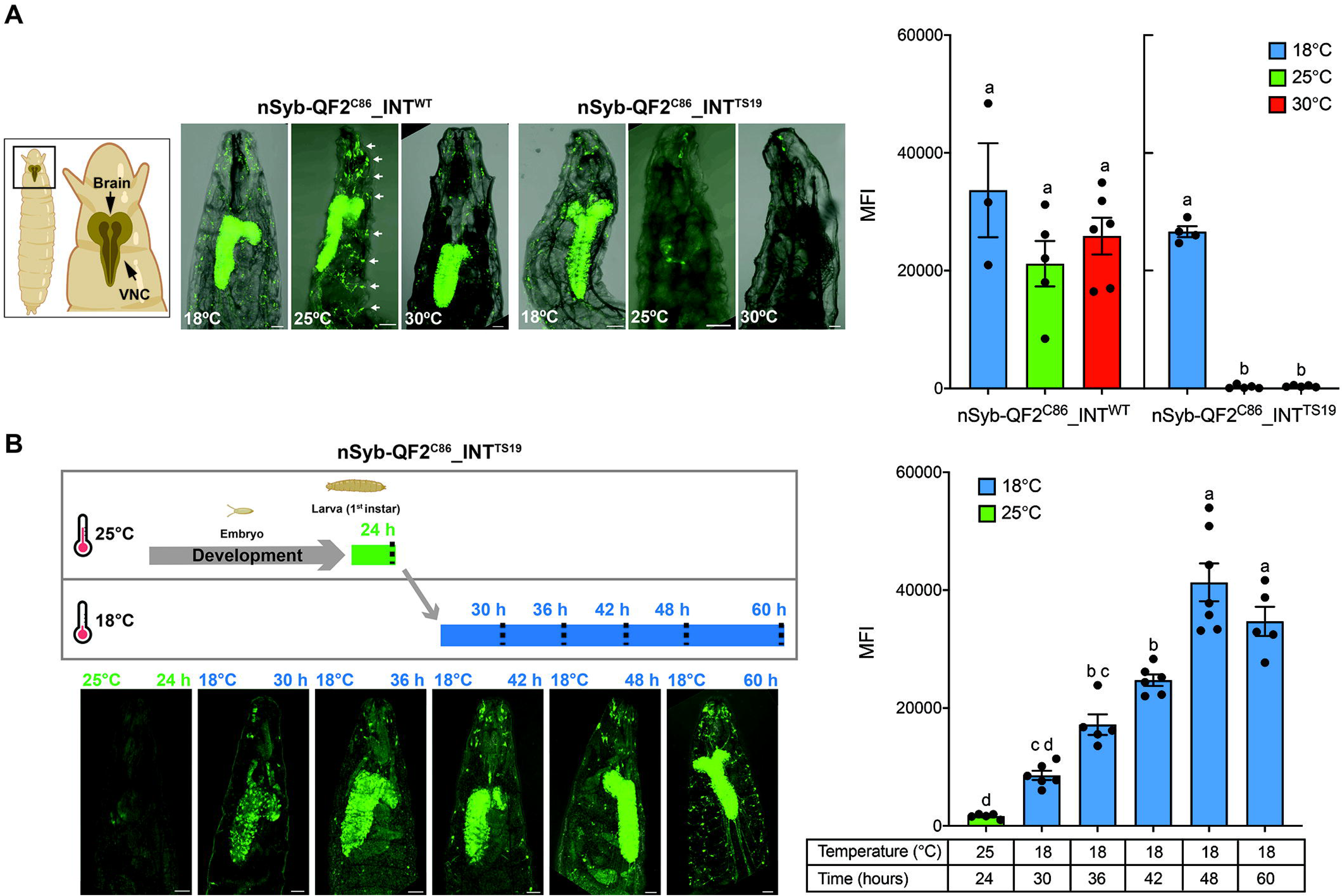
Pan-neuronal expression of GFP in *nSyb-QF2^C86^_INT^TS19^* larvae. **A)** Live confocal images of 1^st^ to 2^nd^ instar larvae containing indicated intein containing *nSyb-QF2* drivers and *10xQUAS-6xGFP* reporter reared at indicated temperature. GFP expression is temperature-dependent under the control of the *nSyb-QF2^C86^_INT^TS19^* driver, while the expression is robust at all temperatures with the *nSyb-QF2_INT^WT^* driver. GFP staining is also clearly visible in the PNS (white arrows). **B)** Upon temperature shift to a permissive temperature for 24 hours, GFP expression is fully restored under the control *nSyb-QF2^C86^_INT^TS19^* driver. Live GFP imaging was performed on whole-mount preparations from 1^st^ to 2^nd^ instar larvae. Scale bars are 50 µm. GFP expression was quantified and displayed as MFI (mean fluorescence intensity) in the histogram. Each bar represents the mean ± SEM of MFI (n = 3-7 MFI measurements in the larval CNS). Dots indicate MFI measured from individual CNS of larva. Ordinary one-way ANOVA with Tukey’s multiple comparison tests (p < 0.05). Different letters indicate statistically significant differences.

### Temporal control of neuronal activity using Kir2.1

A critical feature of the standard Q-system is a suppressor component (QS) provided via a third transgene that is controlled by a constitutively active promoter. Suppression of QF by QS is essential for functional analyses of cells and tissues during late larval development or adult flies driving QUAS reporters that silence neurons, such as the inward rectifying potassium channel Kir2.1 (Baines et al. 2001) and tetanus toxin (Sweeney et al. 1995), or encode cell death genes, such as reaper (White et al. 1994, 1996). To functionally establish whether QF2^C86^_INT^ts^ activators controlling reporter expression with systemic lethal effects can replace the need of a suppressor, we first examined OSN mediated olfactory preference behavior in flies conditionally expressing *QUAS-Kir2.1*. It is well established that a subset of *Orco-* expressing neurons and ORCO containing olfactory receptor complexes are critical for responding to ethyl acetate (Steck et al. 2012; Mansourian and Stensmyr 2015; Kepchia et al. 2017), which we confirmed using a T-maze assay (Figure 4A). We therefore assessed the behavioral impact of Kir2.1 expression in *Orco* neurons driven either by standard QF2 or by intein-regulated QF2 constructs (*Orco- QF2^C86^_INT^WT^, Orco-QF2^C86^_INT^J1^*) in flies kept at 25°C and 30°C. Control flies (reporter or driver alone flies) exhibited strong attraction to ethyl acetate vs solvent, while *Orco-QF2/QUAS-Kir2.1* and *Orco-QF2^C86^_INT^WT^/QUAS-Kir2.1* flies showed no preference, regardless of temperature (Figure 4B). In contrast, *Orco-QF2^C86^_INT^J1^/QUAS-Kir2.1* flies showed preference for ethyl acetate when kept at 30°C but were not attracted to it when kept at 25°C (Figure 4B). Furthermore, *Drosophila* raised at 30°C throughout development until two days after eclosion and downshifted to 25° C lost ethyl acetate preference within 1 days, indicating that intein splicing in QF2^C86^_INT^J1^, and hence Kir2.1 mediated neuronal silencing, is induced within 24 hours (Figure 4C).

**Figure 4.**
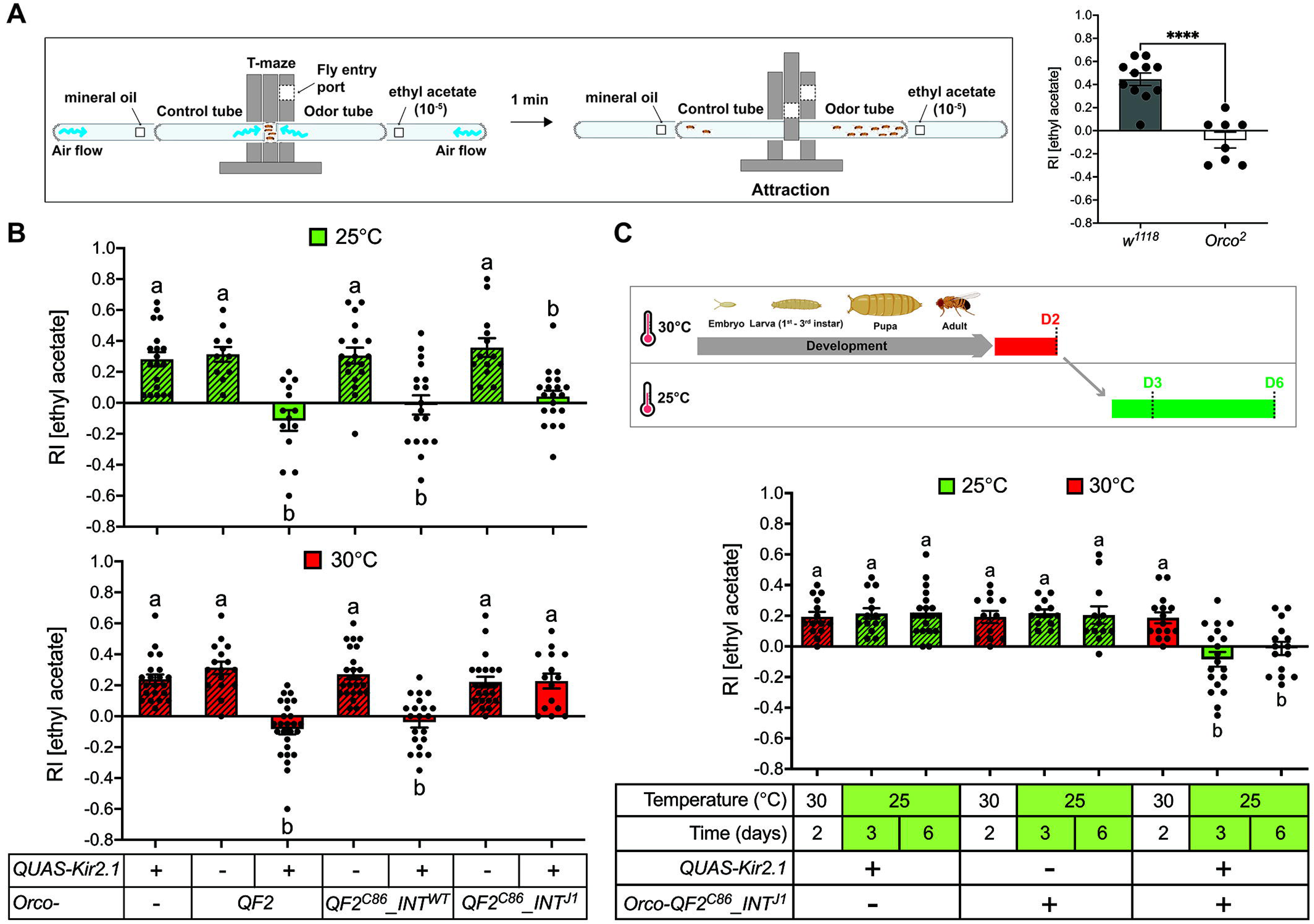
Kir2.1-mediated inhibition by Orco-QF2^C86^_INT^J1^ results in loss of odor preference. **A)** *Orco* mutants show loss of odor preference in T-maze assay. Experimental set up (diagram) uses an “elevator” to slide 20 flies into the center of the T-maze. Ethyl acetate containing air and control flow towards the flies, which are given one minute to choose either the odor or the control tube. Filter paper soaked with 50 µl of ethyl acetate (10^-5^-fold diluted in mineral oil) and solvent (pure mineral oil), respectively, are placed in a tube attached to the odor or the control tube. All tubes are perforated at the bottom to allow airflow generated via a vacuum pump. Homozygous *Orco^2^* mutant flies show no attraction to ethyl acetate, while control flies (w^1118^) exhibit clear preference. Asterisks indicate a significant difference between *w^1118^* and *w^1118^;+;Orco^2^* flies (two- tailed, unpaired Welch’s t test, **** p < 0.0001). **B)** *Orco-QF2^C86^_INT^J1^* mediated silencing using Kir2.1 results in loss of ethyl acetate attraction. Flies reared at 25°C display complete loss of ethyl acetate attraction (top graph, green), while flies reared at 30°C exhibit strong preference (bottom graph, red). Each bar represents the mean ± SEM of RI (response index) (n = 11- 24 assays). Dots indicate RI from individual olfactory choice assays of 20 flies. Ordinary one-way ANOVA with Tukey’s multiple comparison tests (p < 0.05). Different letters indicate statistically significant differences. Fly genotypes: controls (green or red pattern): *w^1118^;SP* or *CyO/+;QUAS-Kir2.1/+*, *w^1118^;+;Orco-QF2/+*, *w^1118^;+;Orco-QF2^C86^_INT^WT^/+*, *w^1118^;+;Orco-QF2^C86^_INT^J1^/+*, experimental groups (green or red): *w^1118^;SP or CyO/+;Orco- QF2/QUAS-Kir2.1*, *w^1118^;SP or CyO/+;Orco-QF2^C86^_INT^WT^/QUAS-Kir2.1*, and *w^1118^;SP or CyO/+;Orco-QF2^C86^_INT^J1^/QUAS-Kir2.1*. **C)** *Orco-QF2^C86^_INT^J1^/QUAS-Kir2.1* flies raised at restrictive temperature loose acetyl acetate preference within 24 hours after shifting to permissive temperature. (Top) Experimental temperature-shift paradigm. *Drosophila* was reared at 30°C through development and, after adult eclosion, subjected to temperature shift to a permissive (25°C) condition for the indicated durations. Arrows indicate transitions between temperatures. (Bottom) Each bar represents the mean ± SEM of RI (response index) (n = 12 - 20 assays). Each dot indicates RI from individual olfactory choice assay of 20 flies. Ordinary one-way ANOVA with Tukey’s multiple comparison tests (p < 0.05). Different letters indicate statistically significant differences. Fly genotypes: controls (green or red pattern): *w^1118^;SP* or *CyO/+;QUAS-Kir2.1/+* and *w^1118^;+;Orco- QF2^C86^_INT^J1^/+*, experimental group (green or red): *w^1118^;SP or CyO/+;Orco- QF2^C86^_INT^J1^/QUAS-Kir2.1*.

Next, we characterized development and adult flies conditionally expressing *UAS-Kir2.1* under the control of *nSyb-QF2^C86^_INT^TS19^* (Figure 5). Expression of Kir2.1 in embryos at permissive temperature resulted in embryonic lethality, as did expression under control of a constitutive *nSyb-QF2* driver, consistent with a previous report (Nitabach et al. 2002), whereas keeping *nSyb-QF2^C86^_INT^TS19^/QUAS-Kir2.1* embryos and larvae at restrictive temperature had no effect on survival and resulted in eclosion of flies at rates indistinguishable from animals lacking either driver (Figure 5A). To further explore the suitability of the Q^INT^ system for conditional expression, we examined the effects of loss of neuronal activity in adult flies. We tested *nSyb- QF2^C86^_INT^TS19^/SP or CyO;QUAS-Kir2.1/+* flies raised at 25 °C in a standard motor behavior assay (geotaxis assay) after downshifting them to 18 °C for 1, 3, 4 and 5 days (Figure 5B, top panels). This assay measures the natural negative geotaxis, a behavior that can be quantified by measuring climbing activity after flies are tapped to the bottom of a vial (Figure 5B)(Bainton et al. 2000). Control flies lacking *QUAS-Kir2.1* (Figure 5B) performed well, with 86 - 96% of flies reaching the 8 cm mark within 10 seconds across all five days, indicating that their climbing activity (and hence neural coordination) is intact. Experimental flies (*nSyb-QF2^C86^_INT^TS19^/SP or CyO;QUAS-Kir2.1/+*) performed equally well at 25 °C, but their climbing ability deteriorates severely within 3 days after downshift to 18 °C (34% in males and 18% in females; Figure 5B), leading to complete paralysis after 5 days. Altogether, our studies show that intein-based conditional QF2 expression allows for temporal and reversible control of protein function, thereby eliminating the need of the drug-dependent suppressor QS.

**Figure 5.**
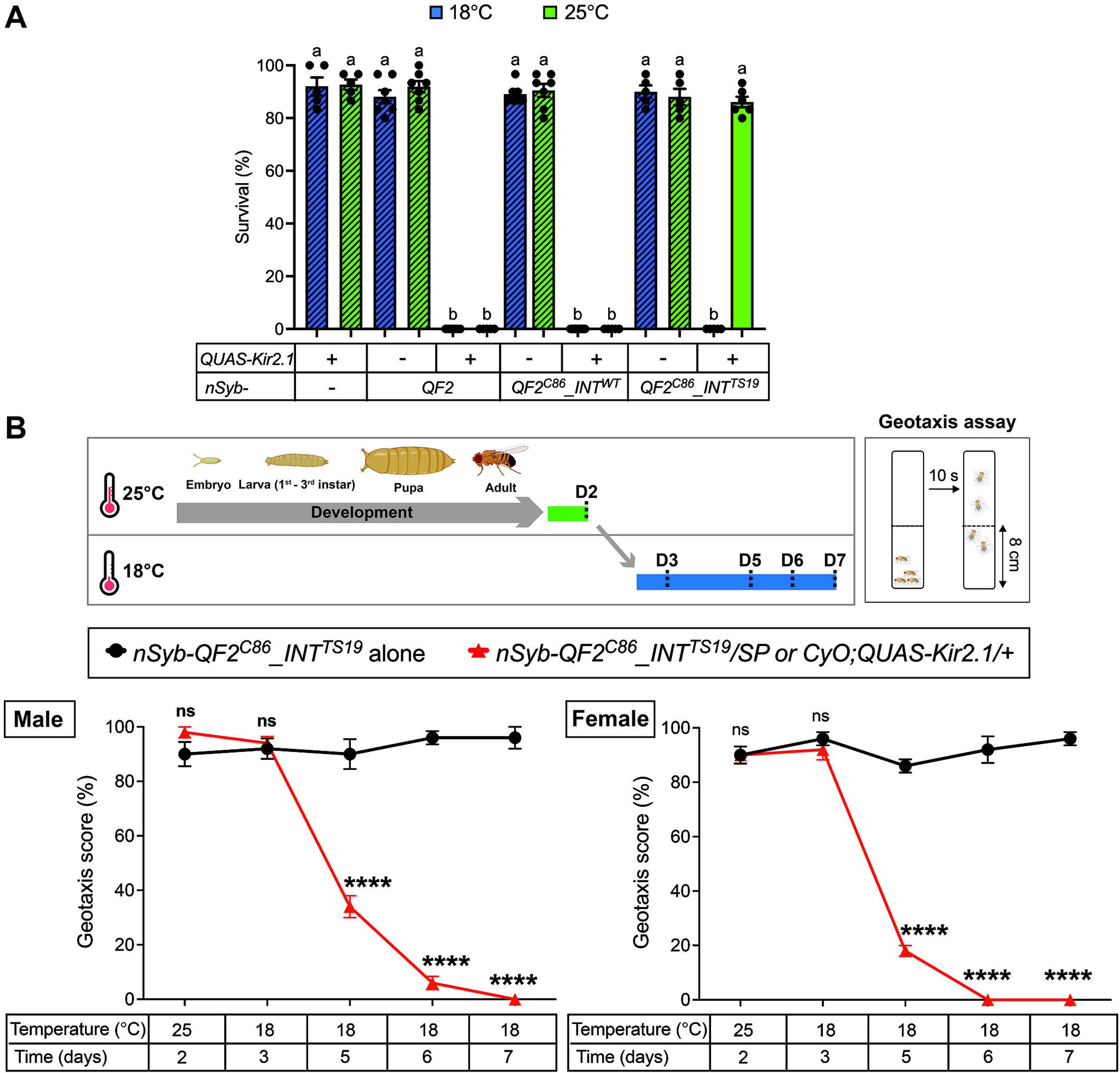
Conditional pan-neuronal silencing during and after development. **A)** Embryos develop normally and reach adulthood at restrictive temperature. *nSyb- QF2^C86^_INT^TS19^/SP or CyO; QUAS-Kir2.1/+* animals raised at restrictive temperature (green bar) develop normally and hatch at rates indistinguishable to controls (driver only and reporter only animals; hatched bars), while the same animals kept at permissive temperature die during embryogenesis as do animals kept at all temperatures with standard *nSyb-QF2* or *nSyb- QF2^C86^_INT^WT^* drivers. Survival (%) = (N_eclosed flies_ / N_eggs_) x 100. N indicates the number of eclosed flies and eggs, respectively. Each bar represents the mean ± SEM of survival (n = 5- 11 experiments). Each dot indicates individual survival experiment. Ordinary one-way ANOVA with Tukey’s multiple comparison tests (p < 0.05). Different letters indicate statistically significant differences. Fly genotypes: controls (blue or green pattern): *w^1118^;SP* or *CyO/+;QUAS-Kir2.1/+*, *w^1118^;+;nSyb-QF2/+*, *w^1118^;nSyb-QF2^C86^_INT^WT^/+,* and *w^1118^;nSyb-QF2^C86^_INT^TS19^/+*, experimental groups (blue or green): *w^1118^;SP or CyO/+;nSyb-QF2/QUAS-Kir2.1*, *w^1118^;SP or CyO/ nSyb-QF2^C86^_INT^WT^;QUAS-Kir2.1/+* and *w^1118^;SP or CyO/ nSyb-QF2^C86^_INT^TS19^;QUAS- Kir2.1/+*. **B)** Kir2.1-mediated neuronal silencing leads to loss of motor activity and paralysis in adult flies shifted to permissive temperature. Top panel shows experimental strategy for geotaxis assay. 10 experimental (*w^1118^;SP or CyO/nSyb-QF2^C86^_INT^TS19^;QUAS-Kir2.1/+*) or control flies (*w^1118^;nSyb- QF2^C86^_INT^TS19^/+*) raised at restrictive (25°C) temperature were tested prior to and 3, 5, 6 and 7 days after downshift to permissive (18°C) temperature for negative geotaxis, by gently tapping them to the bottom of the vial and recording the number of flies reaching the threshold line within 10 seconds. Males and females were analyzed separately and showed no obvious difference in geotaxis. Geotaxis score is percentage of flies above 8 cm within 10 s. Each dot or triangle represents the mean ± SEM of geotaxis score (n = 5 assays). Asterisks indicate a significant difference between *w^1118^;SP or CyO/nSyb-QF2^C86^_INT^TS19^;QUAS-Kir2.1/+* and *w^1118^;nSyb- QF2^C86^_INT^TS19^/+* flies (Two-tailed, unpaired Welch’s t test, **** p < 0.0001, ns: not significant).

## Discussion

Powerful binary expression systems have long been limited to classic genetic model systems, such as *C. elegans*, *D. melanogaster and M. musculus*. With the advancement of genome editing, such as *Crispr/Cas9*, these expression systems are primed to be used more widely in other animals as analytical and functional tools examining gene expression and function at the cellular level. A key feature of these systems is their reliance on the regulatory elements of the gene promoter driving the transcriptional activator, which imposes tissue-specific, developmental and/or physiological control, providing investigators access to specific cell types and organs, or tissues during development. If deployed under the control of ubiquitous promoters or promoters expressed across a wide cell lineage (such as neurons), a key advantage comes along with a major drawback, namely the lack of temporal control. This caveat has been counteracted using expression systems regulatable by chemicals, such as tetracycline and doxycycline in the Tet on/off system and quinic acid and QS in the Q-system. However, temporal control of QF activity is complicated by the fact it relies on QS as a third genetic component, whose functional removal requires dietary supplementation of quinic acid, which tastes bitter and is poorly consumed at concentrations high enough to be effective, especially for genes expressed in the CNS.

For applications in Drosophila, the Q^INT^-system presented in this paper simplifies current binary expression systems by providing temporal and reversible gene expression control without the need of a tertiary suppressor component (QS and GAL80, respectively) and simultaneously eliminates the caveats associated with dietary supplementation of quinic acid. Activation of QUAS reporters in the Q^INT^-system is efficiently achieved in the CNS within 24 hours (Figure 3) and OSNs within 4 days (Figure 2) after larvae/flies were shifted from the restrictive to permissive temperature conditions, providing an ideal platform for manipulating functionality of neurons, via neural activators and suppressors (QUAS-CsChrimson, QUAS-Kir2.1), Calcium-dependent sensors (QUAS-GCaMP, QUAS-CaLex), or gene-specific suppressors (QUAS-RNAi constructs). In addition, the HACK system (Lin and Potter 2016) provides a simple platform for converting standard GAL4 and QF drivers into QF2_INT^ts^ drivers.

Based on the temperature profiles, the intein-based QF2 drivers fall largely into four groups. Orco-QF2^C86^_INT^WT^ is constitutively active at all temperature, generating QUAS-GFP expression profiles indistinguishable from the standard Orco-QF2 driver (Figure 1D). The four drivers with temperature-sensitive intein modules are all active at 18 °C and inactive at 30 °C. At 25 °C, only Orco-QF2^C86^_INT^J1^ is active (at about 40% of the level observed in Orco-QF2_INT^WT^), while Orco-QF2^C86^_INT^F1^ and Orco-QF2^C86^_INT^TS19^ fail to drive even sparse expression of the QUAS-GFP reporter (Figure 1D). Finally, Orco-QF2^C86^_INT^J2^ exhibits sparce expression at 25 °C, suggesting that its threshold temperature is likely somewhere between 18 °C and 25 °C. It is important to recognize that none of these temperature-sensitive VMA1 intein modules have a sharp threshold point, but transition from (almost) fully spliced to none-spliced over a small centigrade range, as previously established in yeast (Zeidler et al. 2004; Tan et al. 2009). Regardless, investigators can choose between the four modules that best suits their experimental questions. Moreover, Q55 in the VMA1 intein protein might serve as a prime site for additional mutations with a higher temperature shut off point, given that QF2^C86^_INT^J1^ (Q55P) provides robust activity at 25 °C. Such drivers may be especially desirable when QF2^C86^_INT^ts^ drivers are introduced into other insects, such as A. aegypti, which thrive best between 25 °C and 29 °C, and can tolerate temperatures relatively well up to 33 °C.

While the GAL4 system is more widely used in *Drosophila* than the Q-system, its usefulness in other models has been limited due to toxicity and epigenetic silencing of the reporter gene (Goll et al. 2009; Akitake et al. 2011; Coutinho-Abreu and Akbari 2023). For these reasons, the Q-system has been introduced into numerous other insects as well as poikilotherm vertebrates (*D. rerio*) with modified transcriptional activators, such as QF2 and QF2^w^ which show little or no toxicity. Moreover, the QF2 binding site (QUAS) is not prone to genome silencing unlike the GAL4 binding site (UAS) as it is devoid of CpG dinucleotides. Thus, QF2_INT^ts^ drivers will provide new and precise command in functional analyses of genes at specific times in development or in adult animals by enabling temporal activation. Temporal control of broadly expressed QF2 drivers is essential when studying gene and cell function by means of RNAi or expression of cell death genes, respectively, or by inducing and suppressing neural activity using optogenetically stimulated neural activators and Kir2.1, respectively (Liang et al. 2013; Simpson and Looger 2018; DeAngelis et al. 2020; Yawo et al. 2021). Temperature-sensitivity is only one of several regulatable parameters that has been levied on inteins, and thus, it is likely that other control mechanisms, such as photogenic stimuli, can be exploited to control intein splicing from QF2 and other transactivators in the future, which would allow expansion of temporal control to warm-blooded vertebrates.

Incorporation of the same INT^ts^ variants into different proteins (GAL4, GAL80 and QF2) revealed new insights into the role of host protein context on temperature sensitivity of specific INT modules: While INT^F1^ exhibits a similar temperature profiles in splicing from both GAL4 (expressed in *S. cerevisiae* ; Tan et al. 2009) and QF2 (*D. melanogaster*; this report) with efficient splicing at 18° C but little or no splicing at 25°C and above, INT^TS19^ splicing from GAL4 (*S. cerevisiae* ; Tan et al. 2009) appears less sensitive to increasing temperature compared to GAL80 or QF2 (both *D. melanogaster*). Specifically, virtually no loss in splicing efficiency from GAL4 is observed between temperatures ranging from 18 °C to 25 °C, with still high level of splicing observed even at 28 °C, whereas splicing from GAL80 and QF2 is severely diminished or completely lost (Supplementary Table 1). Thus, temperature sensitivity is dependent not only on a particular mutation within the intein but is also affected by the host protein sequence context.

Lastly, use of INT^ts^ modules presented here might be exploited in several other contexts. For example, they might be inserted in the coding sequence of genes, for which temperature sensitive alleles would be desirable, such as genes with diverse essential functions during development, greatly facilitating functional analysis in adult flies. Finally, the vast collection of standard QF and QF2 drivers would benefit from a QS suppressor that does not rely on the difficult to administer quinic acid, which should be straightforward by generating temperature sensitive intein-containing suppressor (QS_INT^ts^).

## Material & Methods

### Key Resources Table

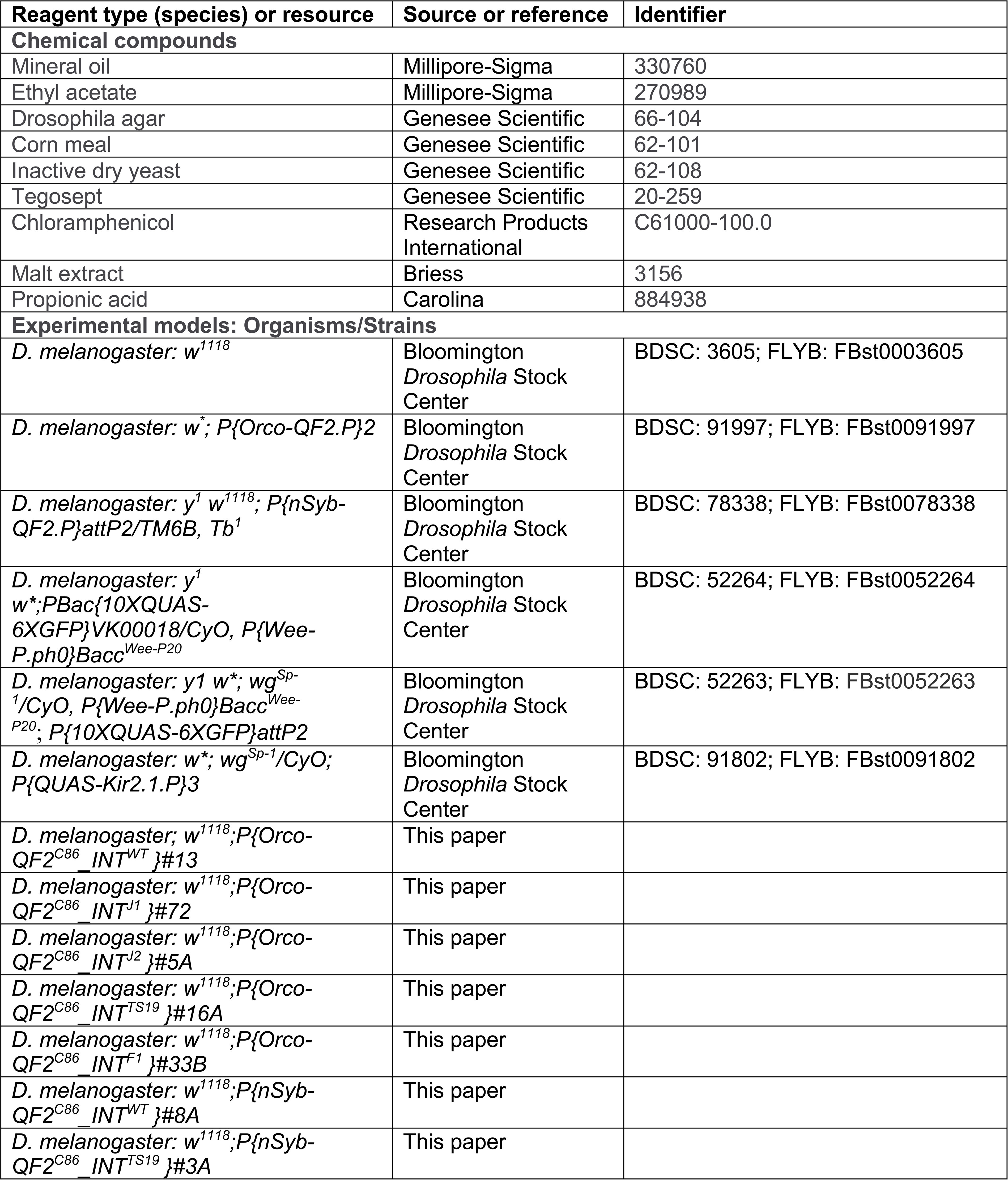
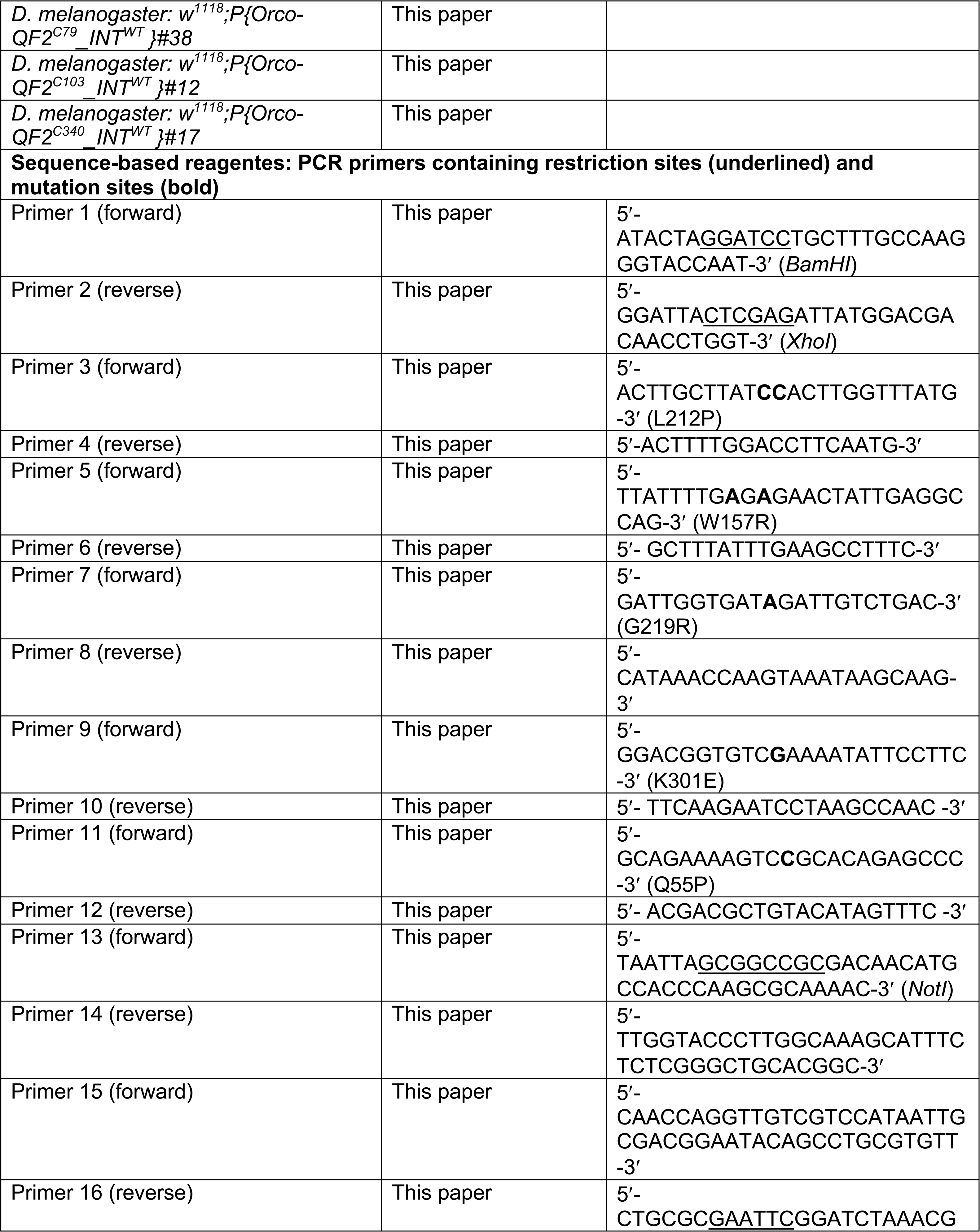
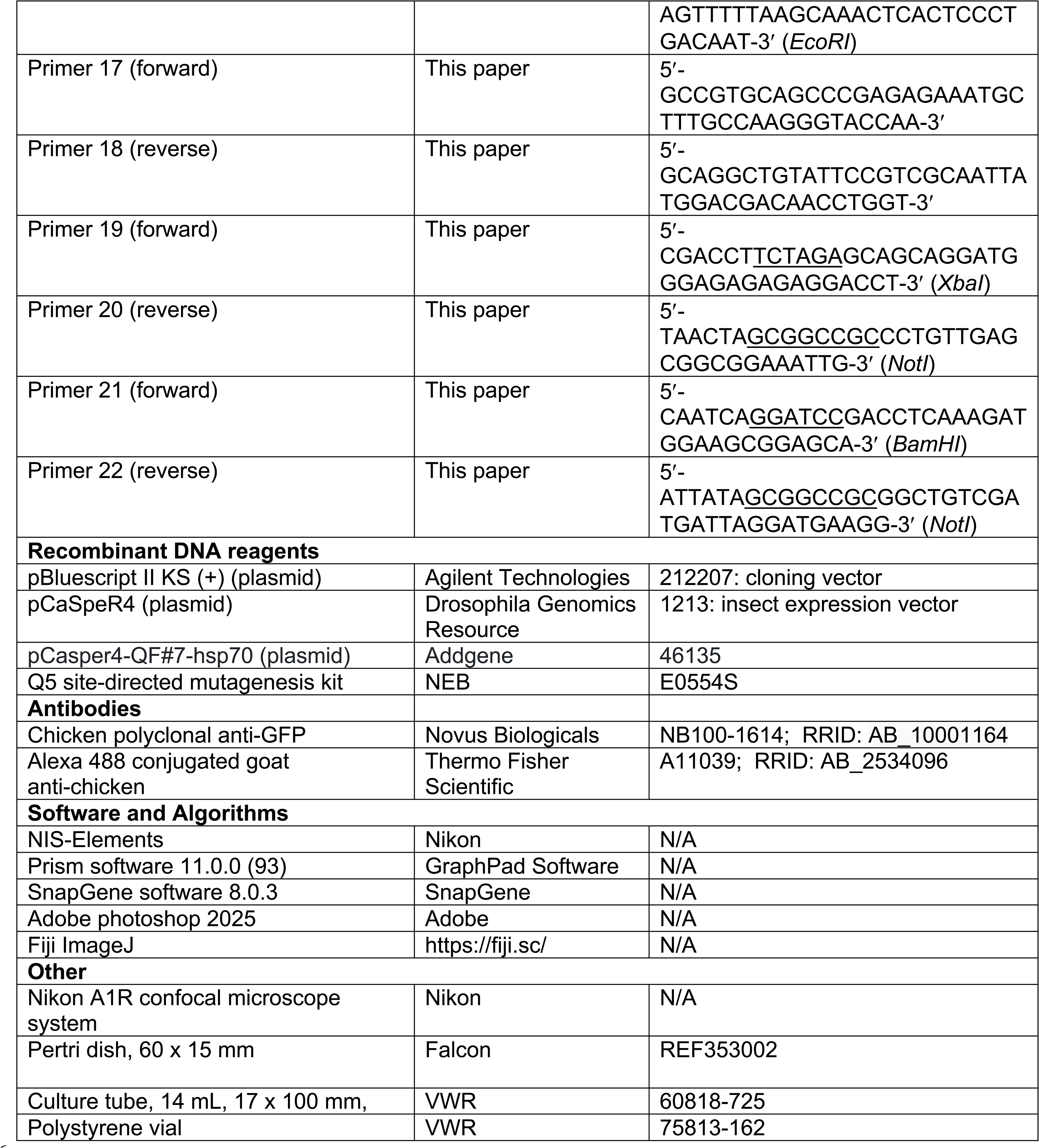

### Molecular Biology

All cloning was performed using pCaSpeR4 (Drosophila Genomics Resource Center, Bloomington, IN, stock number 1213) and pBluescript II KS (+) (referred to as pBS henceforth) (Agilent Technologies, Santa Clara, CA, Cat No. 212207) plasmid vectors.

#### pBS-intein^WT^

To amplify the vacuolar membrane H+-ATPase wild type intein (*intein^WT^*), PCR (98°C for 10 sec, 65°C for 30 sec, 72°C for 1 min for 35 cycles) was performed using Primer 1, 2 (see Methods) and *Saccharomyces cerevisiae* genomic DNA (provided by Dr. Michael Polymenis) as a template. The 1362 bp of PCR fragment was subcloned using *BamHI* and *XhoI* sites of pBS and the insert sequence of pBS-intein^WT^ was confirmed by DNA sequence analysis.

#### pBS-QF2^C86^-Intein^WT^

*Intein^WT^* module was inserted immediately 5¢ to C86 of the *QF2* gene using overlap extension PCR (Hilgarth and Lanigan 2020), generating QF2^C86^-Intein^WT^ fusion, under the following conditions: 98°C for 10 sec, 72°C for 1 min for 35 cycles. pCasper4-QF#7-hsp70 (Addgene plasmid # 46135; Riabinina et al. 2015) served as template to amplify the N- (M1 to K85) and C- (C86 to Q350) terminal QF2 using primers 13 and 14 (295 bp) and 15 and 16 (1096 bp), respectively. The Intein^WT^ module (1402 bp) was amplified from pBS-Intein^WT^ with primers 17 and 18. Equal micromolar amounts of purified PCR products (QIAquick PCR Purification kit, QIAGEN) were mixed and subjected to the secondary PCR (98°C for 10 sec, 72°C for 1 min 30 sec for 35 cycles) with primers 13 and 16. The assembled 2710 bp fragments were digested with *NotI* and *EcoRI*, gel purified and subcloned into *NotI* and *EcoRI* digested pBS, generating pBS- QF2^C86^-Intein^WT^. The entire insert was sequenced for verification.

#### Intein^TS^ modules

Temperature-sensitive inteins were obtained by site-specific mutagenesis (Q5 site-directed mutagenesis kit, NEB Cat# E0554S) and standard PCR (98°C for 10 sec, 56 ∼ 64°C for 30 sec, 72°C for 3 min for 25 cycles) using plasmid DNA as a template. All cloning was conducted in the pBS vector.

#### *Intein^J2^* and *Intein^J1^*

*Intein^J2^* (K301E) and *Intein^J1^* (Q55P) were generated by PCR amplification of pBS-QF2^C86^-Intein^WT^ using primers 9 and 10, and 11 and 12, respectively.

#### *Intein^F1^* and *Intein^TS19^*

To obtain *Intein^F1^* (L212P), pBS-Intein^WT^ served as template and was amplified with primers 3 and 4. *Intein^TS19^*, which has two point mutations (W157R and G219R), was obtained in two steps: pBS-Intein^WT^ served as the first template, was amplified with primers 5 and 6 and cloned (see below), yielding pBS-Intein^TS19-1^ (W157R). This new template was then amplified with primers 7 and 8, yielding the double mutant pBS-Intein^TS19^. Amplified fragments, phosphorylated using T4 DNA kinase, circularized using DNA ligase and treated with *DpnI* to remove starting template. DNA was transformed into NEB 5a competent *E. coli* cells, and plasmid were grown and sequenced to confirm introduction of mutations.

*Intein^F1^* and *Intein^TS19^* modules were inserted immediately 5¢ to C86 of the *QF2* gene using overlap extension PCR (Hilgarth and Lanigan 2020), generating pBS-QF2^C86^-Intein^ts^, under the following conditions: 98°C for 10 sec, 72°C for 1 min for 35 cycles. pCasper4-QF#7-hsp70 (Addgene plasmid # 46135; Riabinina et al. 2015) served as template to amplify the N- (M1 to K85) and C- (C86 to Q350) terminal QF2 using primers 13 and 14 (295 bp) and 15 and 16 (1096 bp), respectively. The Intein^F1^ and Intein^TS19^ modules (1402 bp) were amplified from pBS-Intein^F1^ and pBS-Intein^TS19^, respectively, with primers 17 and 18. Equal micromolar amounts of purified PCR products (QIAquick PCR Purification kit, QIAGEN) were mixed and subjected to the secondary PCR (98°C for 10 sec, 72°C for 1 min 30 sec for 35 cycles) with primers 13 and 16. The assembled 2710 bp fragments were digested with *NotI* and *EcoRI*, gel purified and subcloned into *NotI* and *EcoRI* digested pBS, generating pBS-QF2^C86^-Intein^F1^ and pBS-QF2^C86^-Intein^TS19^. Entire inserts were sequenced for verification.

Cloning of inteins 5¢ to C79, C103 and C340 were performed in a similar manner.

#### *Orco* and *nSyb* promotor

The *Orco* promoter (corresponding to 2642 bp upstream of the ATG start codon; Wang et al. 2003) was PCR-amplified from *w^1118^* genomic DNA using Primer 19 and 20, subcloned into pBS using XbaI and NotI sites to generate pBS-Orco and sequenced for verification. The *nSyb* promotor (corresponding to 1905 bp upstream of the ATG start codon; Addgene plasmid # 46107; Riabinina et al. 2015) was PCR-amplified from genomic DNA from *nSyb-QF2* transgenic flies (BDSC #78338) using Primer 21 and 22, subcloned into pBS using BamHI and NotI sites to generate pBS-nSyb and sequenced for verification. PCR was carried out as follows: 98°C for 10 sec, 72°C for 1 min 30 sec for 35 cycles.

#### pCaSpeR4-Orco-QF2^C86^_INT^X^

*QF2_INTs* were excised from pBS-QF2^C86^_Inteins (WT, F1, TS19, J2 or J1) by NotI/EcoRI digestion and subcloned into NotI/EcoRI digested pCaSpeR4, generating pCaSpeR4-QF2^C86^_Intein^ts^. Subsequently, the Orco promoter from XbaI/NotI digested pBS-Orco was subcloned into the each of the five XbaI/NotI digested pCaSpeR4-QF2^C86^_Intein^ts^ plasmids, generating pCaSpeR4-Orco-QF2^C86^_INT ^X^ (X referring to WT, F1, TS19, J2 or J1).

#### pCaSpeR4-nSyb-QF2^C86^-INT ^WT/TS19^

The nSyb promoter from BamHI/NotI digested pBS-nSyb was subcloned into each of the two BamHI/NotI digested pCaSpeR4-QF2^C86^_INT ^WT/TS19^ plasmids, generating pCaSpeR4-nSyb-QF2^C86^_INT ^WT/TS19^.

#### Transgene Mapping

QF2 transgenes were mapped to a chromosome by crossing homozygous transgenic flies to the double the balancer strain w; CyO/Sp; TM2/TM6B. Precise molecular location of transgenes was performed for the ten most used QF2_INT^ts^ drivers using the inverse PCR (iPCR) (Ochman et al. 1988; A. M. Huang et al. 2009) Integration sites are listed in Supplementary Table 2.

### Generation of transgenic fly lines

pCaSpeR4-Orco-QF2^C86^_INT^X^ and pCaSpeR4-nSyb-QF2^C86^_INT^WT/TS19^ transgenenic flies were generated using standard P-element mediated transformation procedure of *w^1118^* embryos (Rainbow transgenic flies Inc., Camarillo, CA).

### Maintenance of larvae and flies

Fly stocks were kept on standard corn meal food (145 g of agar, 1040 g of corn meal, 2200 g of malt extract, 550 g of inactive dry yeast, 1 g of chloramphenicol, 0.31% (v/v) propionic acid and 0.14% (w/v) tegosept contained in 20 L of water) in plastic vials under 12-hour light/dark cycles at 25°C. For experiments, eggs, larvae and flies were maintained at indicated temperatures, according to the description outlined for the specific experiment. A detailed list of fly strains used for this paper is provided in the Key Resource Table.

### Chemicals

A detailed list of chemicals used for this paper is provided in the Key Resource Table.

### Fluorescence imaging and Immunofluorescence

To examine live GFP expression during development, 1^st^ to 2^nd^ instar larvae were soaked in Vectashield (Vector Lab, Cat# H-1200) for 30 min and mounted on a microscope slide. To examine GFP expression in the adult olfactory organs, antennae and maxillary palps were collected and mounted with Vectashield on a microscope slide. Images were obtained using a Nikon A1R confocal microscope system. Adobe Photoshop 2025 was used to analyze images.

Immunofluorescence was performed as previously described (Task et al. 2022) with minor modification. Flies were aged for 4–8 days in standard corn meal food at indicated temperatures in Supplementary Figure 2 before dissection. The antennae were detached from the head, while the maxillary palps were dissected from the labella. Tissues were fixed in fixation buffer (phosphate buffered saline with 4% paraformaldehyde and 0.3% triton X-100) for 40 min at room temperature. Tissues were washed three times in washing buffer (phosphate buffered saline containing 0.3% triton X-100) for 15 min at room temperature and blocked for 1 h at room temperature in blocking buffer (washing buffer with 5% normal goat serum). Tissues were incubated with chicken anti-GFP (1:500 dilution, Novus Biologicals, Cat No. NB100-1614) at 4 °C for 1 day (for the maxillary palp) or 4 days (for the antenna) in blocking buffer. Tissues were washed three times in washing buffer for 15 min, blocked for 1 h in blocking buffer at room temperature and incubated with goat anti–chicken ALEXA 488 (1:200 dilution, Thermo Fisher Scientific, Cat No. A11039) in blocking buffer at 4 °C for 1 day (for the maxillary palp) or 3 days (for the antenna). Tissues were washed three times in washing buffer for 15 min at room temperature and mounted with VectaShield on a microscope slide. Images were obtained using a Nikon A1R confocal microscope system. All incubations and washings were performed under gentle mixing. Adobe Photoshop 2025 was used to analyze images.

To quantify GFP expression, Fiji ImageJ software (Schindelin et al. 2012) was used as described (Shihan et al. 2021; Jain et al. 2023). Sum projections of confocal z-stacks from the antenna, maxillary palp (Figure 1D, 2C and Supplementary Figure 2) and larval CNS (Figure 3) were created to obtain 2D images representing the total integrated intensity of the GFP signal across the entire z-depth. Region of interests (the antenna, maxillary palp and larval CNS) were marked in 2D images and the mean fluorescence intensity (MFI) was measured. The MFI of nonfluorescent area (background) was measured and subtracted from the MFI of a region of interest to correct background effect on an image. Final MFI = MFI of a region of interest – MFI of background: this step was done in Excel and the final MFI was displayed in histograms.

### Development of pCaSpeR4-nSyb-QF2^C86^_INT ^WT/TS19^/QUAS-Kir2.1 embryos and larvae

Equal number (∼30) of males and females were let to mate for 2 days under a 12–hour light/dark cycle at 25°C, transferred to grape juice agar (1%) plates with yeast paste and allowed to lay eggs for 4 hr at room temperature in the dark. Thirty eggs were placed on standard corn meal food in plates (60 × 15 mm, Falcon Cat # REF353002) and cultured at 18 or 25°C.

### Olfactory choice assay

Olfactory choice assay was conducted using T maze apparatus described in (Helfand and Carlson 1989). 2 – 12 days old flies, kept at specified temperatures (see Figure 4) were sorted by sex in groups of 20 under CO_2_ anesthesia, let to recover for 2 days in standard food vials at the same specified temperature. Prior to the assay, flies were starved for 16 to 28 hours (flies kept at 25°C) or 15 to 20 hours (flies kept at 30°C) in empty vials containing a water saturated Kimwipe (Kimberly Clark, Dallas, TX). Assays were initiated by tapping 20 males or females onto the vertical sliding plate of the T maze. Sliding plate was lowered and flies were allowed to choose between an odor or odor less control tube for 1 minute (see below) under constant airflow. Assays were terminated by lifting a vertical sliding plate and recording the numbers of flies in both tubes. Assays were performed in the dark to avoid visual effect. Odor tube was placed alternately on the right and left side of T maze. The response index (RI) was calculated as follows: RI = (N_odor_ – N_control_) / N_total_, whereby N indicates the number of respective flies. Positive values indicate a preference for ethyl acetate while negative values indicate avoidance (repulsion).

50 µl ethyl acetate (10^-5^ diluted in mineral oil) was used as attractive odor source (MilliporeSigma Cat# 270989 and Cat# 330760) and placed on a filter paper (1cm^2^) in a 14 ml culture tubes (17 x 100 mm, VWR Cat# 60818-725) with perforated bottoms that were plugged onto empty culture tubes with perforated bottoms. The opposite control tube contained 50 µl of mineral oil. Odor and control tubes were connected to T maze and air was continuously drawn through it by vacuum for odor delivery.

### Geotaxis assay

Geotaxis assays were carried out as previously described (Leal and Neckameyer 2002; Ali et al. 2011) with minor modifications. Files were separated by sex under CO_2_ anesthesia into groups of 10 one day after eclosion, let to recover for another day at 25°C. and transferred and kept at 18°C for 1, 3, 4 or 5 days before assayed for negative geotaxis ability. The geotaxis apparatus consisted of two connected empty polystyrene vials (VWR Cat# 75813-162), with one vial marked at 8 cm distance from the bottom (see Figure 5B). Flies were loaded into the geotaxis apparatus by gentle tapping and left undisturbed (standing) for 1 minute. Experiments were initiated by gently tapping flies down to the bottom of a vial and terminated by recording the fraction of flies able to climb above the 8-cm mark after 10 seconds.

### Statistical analysis

Statistical analyses were conducted using Prism software 11 (GraphPad Software). Assessment of survival, geotaxis and olfactory choice assay data was analyzed for normal distribution using D’Agostino & Pearson test and Shapiro-Wilk test. For comparison between multiple groups (Figure 1D, 2C, 3, 4B-C, 5A and Supplementary Figure 2), ordinary one-way ANOVA was performed to test for difference of mean. As a post hoc test, Tukey’s multiple comparison tests were employed to compare two specific groups. For comparison between two groups (Figure 4A and 5B), unpaired Welch’s t test with two-tailed P- value was used.

## Acknowledgements

We thank Nadia Gallegos and Pranav Nalam for assistance with live GFP imaging, and Dr. Michael Polymenis for *S. cerevisiae genomic DNA*. This work was supported by grants from the NIH to HA (NIH-R21AI188026 and NIH-1R21AI191589).

## Notes

### Competing Interest Statement

The authors have declared no competing interest.

### Summary of Updates

We appreciate the enthusiastic comments and insightful suggestions of all reviewers who all agreed on the significance of our study, stating that This work represents a significant technical advance in the genetic control of transgene expression in Drosophila, especially in relation to improving the Q-system toolbox (reviewer 1), This paper presents a novel, interesting and valuable contribution to the genetic tool development for model and non-model organisms (reviewer 2), The authors demonstrated, robust, cell type specific QUAS reporter expression in Drosophila olfactory sensory neurons regulated via temperature shifts and rescue of a lethal phenotype under QF/QUAS control (reviewer 3) and This is a well-executed and practically valuable toolkit paper introducing a clever intein-based simplification of the Q-system. The manuscript is well written, clear, and easy to follow.(reviewer 4). Please find enclosed the revised manuscript in which we addressed all issues raised by the reviewers (quantification of reporter expression, antibody staining, time course of turning QF2C86_INTts on and off, etc), as well as small editing issues that were noted by some of the reviewers. We also added a paragraph in the discussion, prompted by a comment of reviewer 1, where we summarize the major advantages of the new QINT system over the classical Q system. Please see point-by-point response to all comments by the reviewers:

