## Supplementary Figures and Tables for "Q^INT^, a novel simplified QF/QUAS expression system with integrated temporal expression control"

**Supplementary Figure 1.**

A

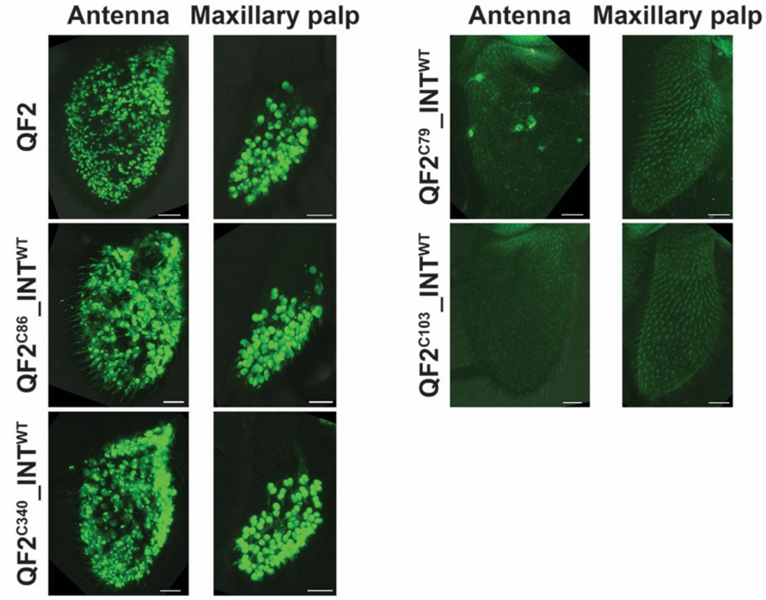


B

| QF2_intein | Intein | Intein insertion | Total number of transgenic lines screened | Total number of transgenic lines driving GFP expression |
|---|---|---|---|---|
| QF2$^{C79}$_INT$^{WT}$ | wild type | C79 | 3 | 0 |
| QF2$^{C86}$_INT$^{WT}$ | wild type | C86 | 4 | 4 |
| QF2$^{C103}$_INT$^{WT}$ | wild type | C103 | 4 | 0 |
| QF2$^{C340}$_INT$^{WT}$ | wild type | C340 | 4 | 4 |

**Expression of *Orco-QF2* containing wild type intein (*INT$^{WT}$*) at upstream of C79, C86, C103, and C340.** Flies homozygous with indicated drivers were crossed to the *10xQUAS-6xGFP* reporter line, and live fluorescence on whole-mount preparations of olfactory organs of adult progeny was performed, with one representative line of each construct shown in A. Scale bars are 20 µm.

(B) Summary of different insertion lines for the wild type intein (*INT$^{WT}$*). Insertion at C79 and C103 within the DNA binding domain resulted in either very poor or no splicing at all, resulting in virtually no GFP expression, while insertion into C86 resulted in highly efficient splicing. Thus, insertion at C86 was used for all temperature sensitive inteins was chosen throughout this study. As shown in Figure 1, insertion of temperature-sensitive inteins (*INT$^{ts}$*) at this site resulted in functional QF2 expression only at permissive temperature. Insertion at C340, residing in the

QF2 activation domain led also to efficient reporter expression, though it cannot be excluded that this insertion might be immune to functional disruption.

**Supplementary Figure 2.**

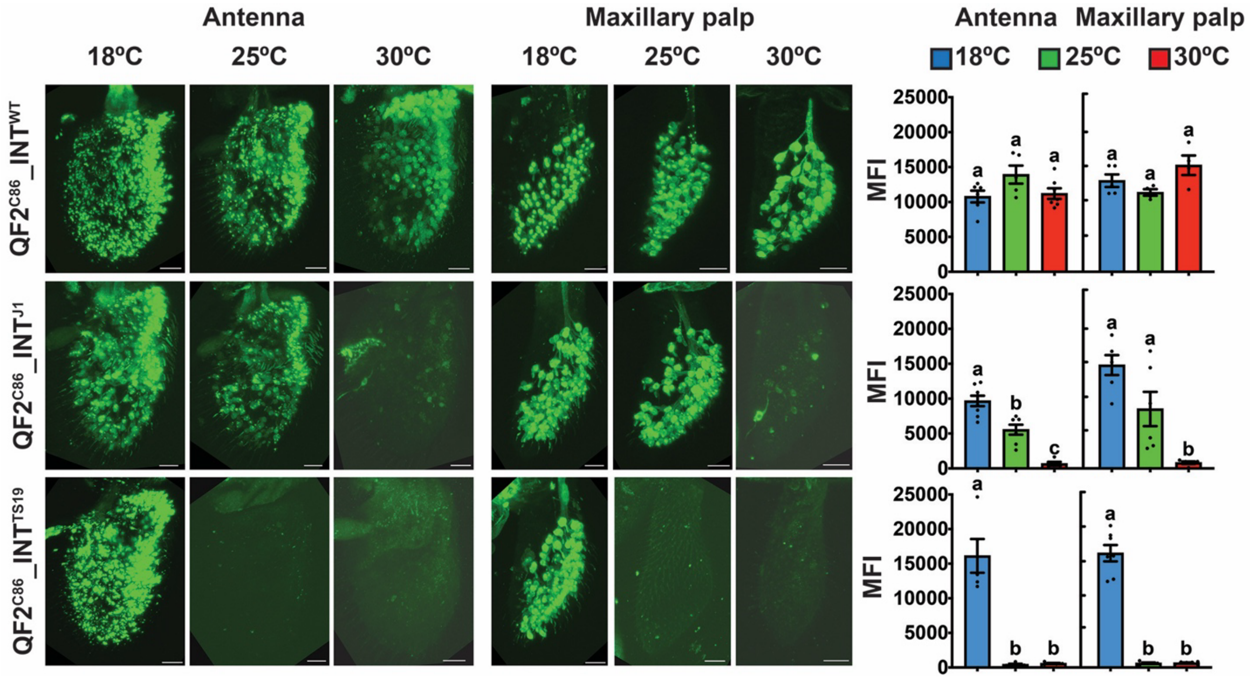


**Immunofluorescence analysis demonstrates robust and reproducible temperature-dependent GFP expression driven by Orco-QF2$^{C86}$_INT in olfactory sensory neurons.** Immunostaining with anti-GFP (green) antibody on whole-mount preparations of the antenna and the maxillary palp from flies of the following genotypes: *w$^{1118}$;10xQUAS-6xGFP/+;Orco-QF2$^{C86}$_INT$^{WT}$/+*, *w$^{1118}$;10xQUAS-6xGFP/+;Orco-QF2$^{C86}$_INT$^{J1}$/+* and *w$^{1118}$;10xQUAS-6xGFP/+;Orco-QF2$^{C86}$_INT$^{TS19}$/+*. GFP expression was quantified and displayed as MFI (mean fluorescence intensity) in the histogram. Each bar represents the mean ± SEM of MFI (n = 4-8 MFI measurements in the antenna/maxillary palp). Dots indicate MFI measured in individual antenna/maxillary palp. Ordinary one-way ANOVA with Tukey's multiple comparison tests ($p < 0.05$). Different letters indicate statistically significant differences. Scale bars are 20 μm.

**Supplementary Table 1. Activity of temperature-sensitive intein (INT$^{ts}$).** Activity of INT$^{ts}$ containing GAL4 activator is indicated as growth of *S. cerevisiae* in galactose medium (Tan et al., 2009). Growth is compared to wild type intein and displayed as no (-), slow (+), accelerated (++), robust (+++) growth. Data from GAL80_INT$^{TS19}$ (Zeidler et al., 2004) and experiments of QF2_INT$^{ts}$ modules are shown in red and green, respectively: +++/+++ indicates full GAL4 suppression/reporter expression; +/+ indicates partial GAL4 suppression/reporter expression; -/- indicates no GAL4 suppression/reporter expression. ND indicates not determined. Intein variant, TS19 was indicated as F19 in Tan et. al (2009).

| Name | Mutation | Intein splicing activity at different temperatures | | | | | | | | | | | |
|---|---|---|---|---|---|---|---|---|---|---|---|---|---|
| | | GAL4 *S. cerevisiae* | | | | | | GAL80 *D. melanogaster* | | | QF2 *D. melanogaster* | | |
| | | 18°C | 22°C | 25°C | 28°C | 30°C | 32°C | 18°C | 25°C | 29°C | 18°C | 25°C | 30°C |
| F1 | L212P | +++ | ++ | + | - | - | - | ND | ND | ND | +++ | - | - |
| TS19 | W157R G219R | +++ | +++ | +++ | ++ | + | - | +++ | + | - | +++ | - | - |
| S40 | Q55P W157R G219R | +++ | +++ | +++ | ++ | + | - | ND | ND | ND | ND | ND | ND |
| S7 | W157R G219R K301E | +++ | + | - | - | - | - | ND | ND | ND | ND | ND | ND |
| J1 | Q55P | ND | ND | ND | ND | ND | ND | ND | ND | ND | +++ | +++ | - |
| J2 | K301E | ND | ND | ND | ND | ND | ND | ND | ND | ND | +++ | + | - |

**Supplementary Table 2. Integration locations of QF2_INT transgenes.**

| Name | Chromosome arm | Genomic location | Cytological location |
|---|---|---|---|
| Orco-QF2$^{C86}$-INT$^{WT}$#13 | 3R | 17794525 | 90C5 |
| Orco-QF2$^{C86}$-INT$^{J1}$#72 | 3L | 699028 | 61C8 |
| Orco-QF2$^{C86}$-INT$^{J2}$#5A | 2R | 16829110 | 53D11 |
| Orco-QF2$^{C86}$-INT$^{TS19}$#16A | 3L | 1638796 | 62A9 |
| Orco-QF2$^{C86}$-INT$^{F1}$#33B | 3L | 14756514 | 70F4 |
| nSyb-QF2$^{C86}$-INT$^{WT}$#8A | 2R | 9578382 | 45F4 |
| nSyb-QF2$^{C86}$-INT$^{TS19}$#3A | 2L | 8522247 | 29D4 |
| Orco-QF2$^{C79}$-INT$^{WT}$#38 | 3R | 5611343 | 83B5 |
| Orco-QF2$^{C103}$-INT$^{WT}$#12 | 2 | Not determined[1] | Not determined[1] |
| Orco-QF2$^{C340}$-INT$^{WT}$#17 | 2R | 22212859 | 58D2 |

[1] Precise integration location of Orco-QF2$^{C103}$_INT$^{WT}$#12 transgene was not determined in the genome due to recovery of repetitive DNA sequence next to insertion site.
